# Characterization of heliorhodopsins detected via functional metagenomics in freshwater Actinobacteria, Chloroflexi and Archaea

**DOI:** 10.1101/2021.02.16.431466

**Authors:** Ariel Chazan, Andrey Rozenberg, Kentaro Mannen, Takashi Nagata, Ran Tahan, Shir Yaish, Shirley Larom, Keiichi Inoue, Oded Béjà, Alina Pushkarev

**Affiliations:** Faculty of Biology, Technion - Israel Institute of Technology, Haifa 32000, Israel; The Institute for Solid State Physics, The University of Tokyo, 5-1-5 Kashiwanoha, Kashiwa, Chiba 277-8581, Japan; PRESTO, Japan Science and Technology Agency, 4-1-8 Honcho, Kawaguchi, Saitama 332-0012, Japan

## Abstract

Rhodopsins are widespread in microbes residing in diverse aquatic environments across the globe. Recently, a new unusual rhodopsin family, the heliorhodopsins (HeRs), was discovered, distributed among diverse bacteria, archaea, eukarya and even viruses. Here, using functional metagenomics on samples from Lake Ha’Hula and Ein Afek reserve, we found and characterized ten HeRs representing divergent members of the family. The expressed HeRs absorb light in the green and yellow wavelengths and originate from Actinobacteria, Chloroflexi and Archaea. The photocycle of the HeR from Chloroflexi revealed a low accumulation of the M-intermediate that we connect to the lack of two conserved histidine residues in transmembrane helices 1 and 2 in this protein. Another of HeR, from Actinobacteria, exhibited an unusually fast photocycle (166 ms, 5 times faster than HeR-48C12). To further explore the still unresolved question of the HeR function, we performed an analysis of protein families among genes neighboring HeRs, in our clones and thousands of other microbes. This analysis revealed a putative connection between HeRs and genes involved in oxidative stress. At the same time, very few protein families were found to distinguish genes surrounding prokaryotic HeRs from those surrounding rhodopsin pumps. The strongest association was found with the DegV family involved in activation of fatty acids and uncharacterized family DUF2177, which allowed us to hypothesize that HeRs are involved in membrane lipid remodeling. This work further establishes functional metagenomics as a simple and fruitful method of screening for new rhodopsins.

**Significance:** The recently discovered divergent rhodopsin family of heliorhodopsins is abundant in freshwater environments. In this study, we sampled a habitat rich in dissolved organic matter to increase our chances of finding spectrally shifted rhodopsins. Using functional metagenomics, diverse heliorhodopsins absorbing green and yellow light were discovered. The metagenomic clones originated from diverse prokaryotic groups: Actinobacteria, Chloroflexi and even Archaea, emphasizing the versatility of the *E. coli* expression system used. Photocycles of representative heliorhodopsins were measured and exhibited diverse kinetic characteristics. Analysis of genes neighboring heliorhodopsins in diverse prokaryotes revealed their putative connection to membrane lipid re-modeling and oxidative stress. Our findings suggest that functional metagenomics is a productive method for the discovery of new and diverse rhodopsins.

## Introduction

Many organisms percept light using rhodopsins (Spudich et al. 2000, Ernst et al. 2014), seven-transmembrane-domain proteins that bind retinal chromophores. Two structurally similar but not directly related rhodopsin families exist: the microbial rhodopsins and the animal rhodopsins (Spudich et al. 2000, Kandori 2020). Found in various microorganisms and even viruses, microbial rhodopsins demonstrate diverse functions that range from generation of ion gradients to harness the energy of light to triggering cell signaling cascades in a light-dependent manner (Kandori 2020).

There exist two different approaches to characterization of novel rhodopsins: (*i*) a top-down approach relying on homology searches in genomic and metagenomic assemblies with subsequent heterologous expression; and (*ii*) a bottom-up approach based on functional metagenomics in which environmental DNA clones are screened for the desired phenotypes, such as pigmentation or light-driven ion-pumping activity (Martinez et al. 2007, Pushkarev and Béjà 2016, Pushkarev et al. 2018). The success of functional genomics is demonstrated by the unexpected discovery of the heliorhodopsins (HeRs) (Pushkarev et al. 2018), a divergent family of microbial rhodopsins with an inverted membrane topology with respect other microbial and animal rhodopsins, that was based on retinal-dependent coloration of a single fosmid clone from Lake Kinneret (Sea of Galilee). Bioinformatic analyses demonstrated that HeRs are abundant and distributed globally, found in archaea, bacteria, eukarya and giant viruses (Pushkarev et al. 2018, Flores-Uribe et al. 2019, Shibukawa et al. 2019, Shihoya et al. 2019). The function of the HeRs remains unknown. Their relatively long photocycle is suggestive of a light-sensing activity (Pushkarev et al. 2018, Shihoya et al. 2019), but support has been also provided for hypotheses of a transporting or an enzymatic activity (Flores-Uribe et al. 2019, Kovalev et al. 2020). HeR genes are found in psychrophiles, mesophiles and even hyperthermophiles, originate from soil, freshwater, marine and hypersaline environments, but are generally lacking from the classical diderm bacteria (Flores-Uribe et al. 2019, Shibukawa et al. 2019).

While HeRs are abundant and diverse, only five proteins have been fully characterized: the actinobacterial HeR 48C12 (Pushkarev et al. 2018), the chloroflexal *Bc*HeR (Shibukawa et al. 2019), the archaeal *T*aHeR, the eukaryotic *Mc*HeR, and the viral *Eh*VHeR (Shihoya et al. 2019). Recently, several additional HeRs were expressed for spectral characterization (Kim et al. 2021).

Since many rhodopsin genes fail to express in *Escherichia coli*, the top-down approach of cloning pre-selected proteins for characterization one by one is a laborious task. Functional metagenomics, on the other hand, screens only for genes that can express in *E. coli*, making a wider phenotype-oriented search for microbial rhodopsins with diverse biophysical properties possible.

When designing the experiment, we were driven by the idea that understanding the mechanism of HeR photoisomerization and ultimately their molecular function would be promoted by finding spectrally-shifted HeRs. We thus searched for a habitat that would favor proteins responding to light with longer wavelengths. In particular, organic matter (OM) has a known direct effect on the absorbed/scattered light in water, with high OM concentrations resulting in almost complete removal of short wavelengths (blue-green) (Stomp et al. 2007, Holtrop et al. 2021). Our choice of sampling sites for this exploration thus fell on two water bodies in northern Israel: a peat lake in the Hula valley and a freshwater pond in the Ein Afek reserve (*SI Appendix*, Fig. S1). Lake Ha’Hula is a re-flooded wetland characterized as a peat lake with high concentration of dissolved substances (Hambright and Zohary 1998). The lake is a pivotal migration station for birds and is known for its highly diverse aquatic biota (Hambright and Zohary 1998). Ein Afek reserve is an array of pools with freshwater fish ponds characteristics, one of the last remnants of the swamp landscapes in northern Israel (Barinova and Romanov 2015).

In this work, we prepared environmental libraries from these freshwater habitats and by using functional screens, we detected ten different HeRs from Actinobacteria, Chloroflexi, and Archaea. As anticipated, one of the HeRs turned out to be the most spectrally-shifted HeR known thus far, with an absorption peak at 562 nm. Concomitantly, it stands out in lacking two conserved key histidine residues in transmembrane helices 1 and 2 which are considered important for HeR photocycle.

In addition to the biophysical characterization of the environmental HeRs, we attempted to approach the still unresolved question of the molecular function of HeRs. Environmental clones (containing ∼40kbp from random genomes) provide some genetic background for the corresponding rhodopsin genes, which gave us the idea of focusing on HeR neighbors as a window to their potential functional partners. We supplemented our small collection with thousands of diverse prokaryotic HeRs from metagenomic and genomic assemblies. This analysis revealed that HeR genes often reside next to genes involved in oxidative stress response, translation and light-dependent proteins, yet the same protein families were found to neighbor rhodopsin protein pumps from the same taxonomic groups. At the same time, the two most characteristic protein families among HeR neighbors appeared to be DegV, a family of fatty-acid binding proteins and an uncharacterized family DUF2177. Based on this evidence, and the ability of HeRs to bind fatty acids, we speculate that HeRs perform a light-dependent enzymatic activity involved in membrane lipid remodeling.

## Results and Discussion

The first representative of the HeR family was discovered and characterized by us from a single clone from Lake Kinneret as part of a functional metagenomic screen (Pushkarev et al. 2018). Since functional metagenomics proved useful in detecting novel rhodopsins (Pushkarev and Béjà 2016, Pushkarev et al. 2018), we deployed our screening method to detect diverse HeR genes from different environmental samples.

In this work, we functionally screened 12,672 environmental clones from Ha’Hula peat lake, and 6336 clones from the Ein Afek reserve. We detected eleven contigs harboring ten distinct HeR genes (Figure 1A; HULAa55C9 and HULAa32G3 being identical) from three different environmental samples (*SI Appendix*, Fig. S1). The isolated environmental clones represented three prokaryotic groups most frequently possessing HeR genes: 9 out of the 11 fosmids originated from Actinobacteria, while the remaining clones HULAa30F3 and HULAa36F11 came from Chloroflexi and Archaea (Thermoplasmata), respectively (Figure 1B and *SI Appendix*, Figs. S2-S4). Detecting an archaeal HeR gene using an *E. coli*-based screen system was surprising as it is believed that the *E. coli* machinery would not recognize native archaeal promoters (Gehring et al. 2016). While archaeal HeRs can be expressed in *E. coli* after codon optimization and changing their promoter and Shine Dalgarno sequences (Shihoya et al. 2019), the archaeal HeR in this study was expressed from its native transcriptional and translational signals.

**Figure 1.**
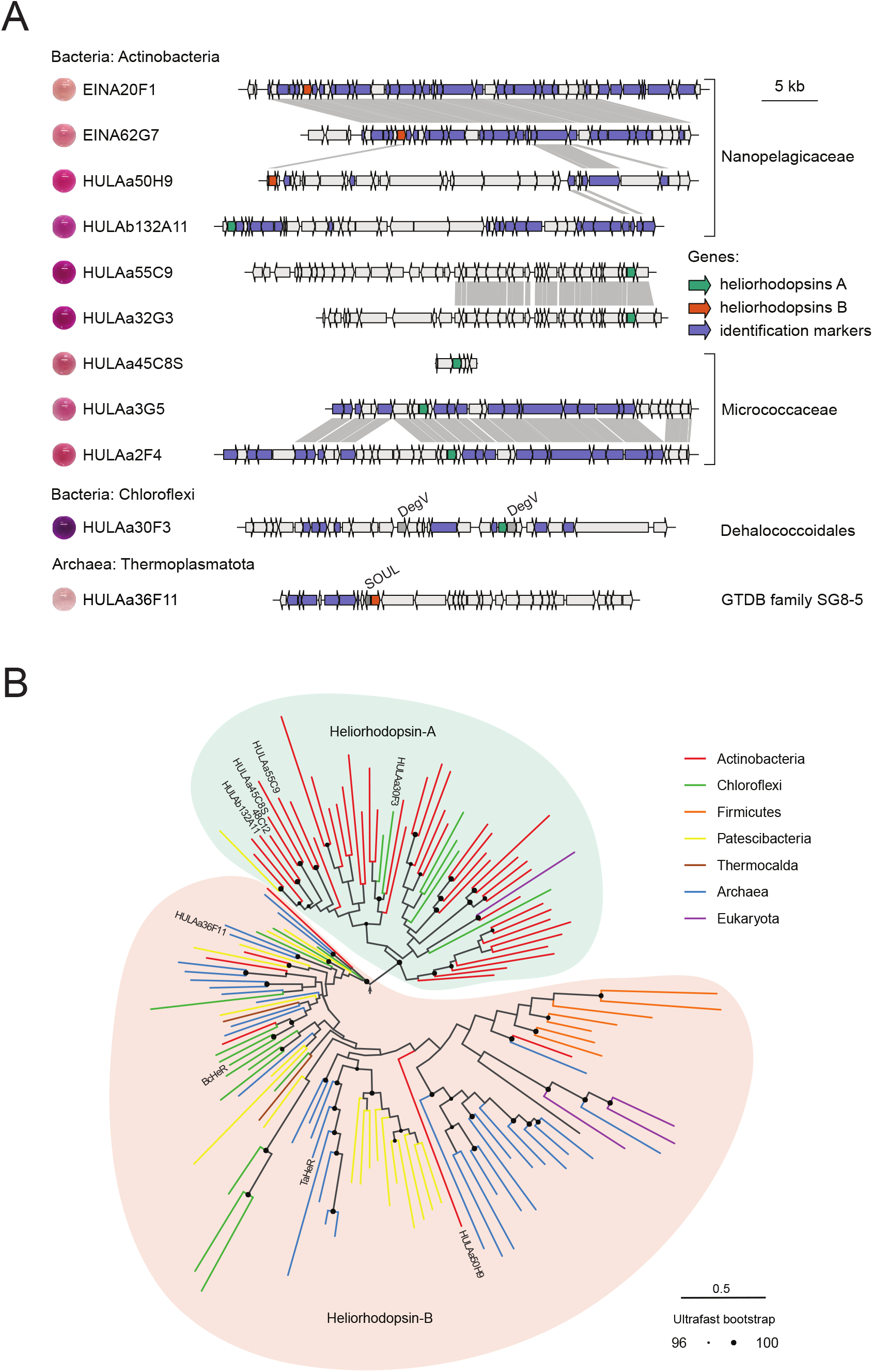
Diversity of prokaryotic HeRs. **A**. Physical map and synteny between the contigs. The HeR genes are indicated in orange and green. Highlighted in blue are genes used for a more detailed taxonomic identification of the fosmids (see *SI Appendix*, Figs. S2-S4). Labels are provided for frequent HeR neighbors (see Figure 4). For each protein, pictures of DDM-solubilized proteins in solution are presented. **B**. Phylogenetic tree of prokaryotic HeRs. The tree is based on 60%-identity clusters of lysine-containing sequences from prokaryotic assemblies and Uniprot after exclusion of the viral and eukaryotic lineages. Clusters that include proteins characterized previously and in this study are labelled correspondingly. The tree is rooted using the minimal ancestor deviation method which places the root at the longest internal branch: between the Actinobacteria-dominated clade, Group A (family 1 in (Kovalev et al. 2020)), and the more diverse Group B. Group assignments are available in *SI Appendix*, Figs. S2-S4.

Interestingly, we found that the inducer concentration optimal for expression of HeR genes varied on a range of several orders of magnitude between different clones when expressed under the same promoter (between 4-2 µl/ml transducer and down to 0.064 µl/ml) (Figure 2A). This variation may be due to differences in codon usage. As previously established (Pushkarev et al. 2018, Shibukawa et al. 2019, Shihoya et al. 2019, Kovalev et al. 2020), none of the detected HeRs showed proton-pumping activity (Figure 2B).

**Figure 2.**
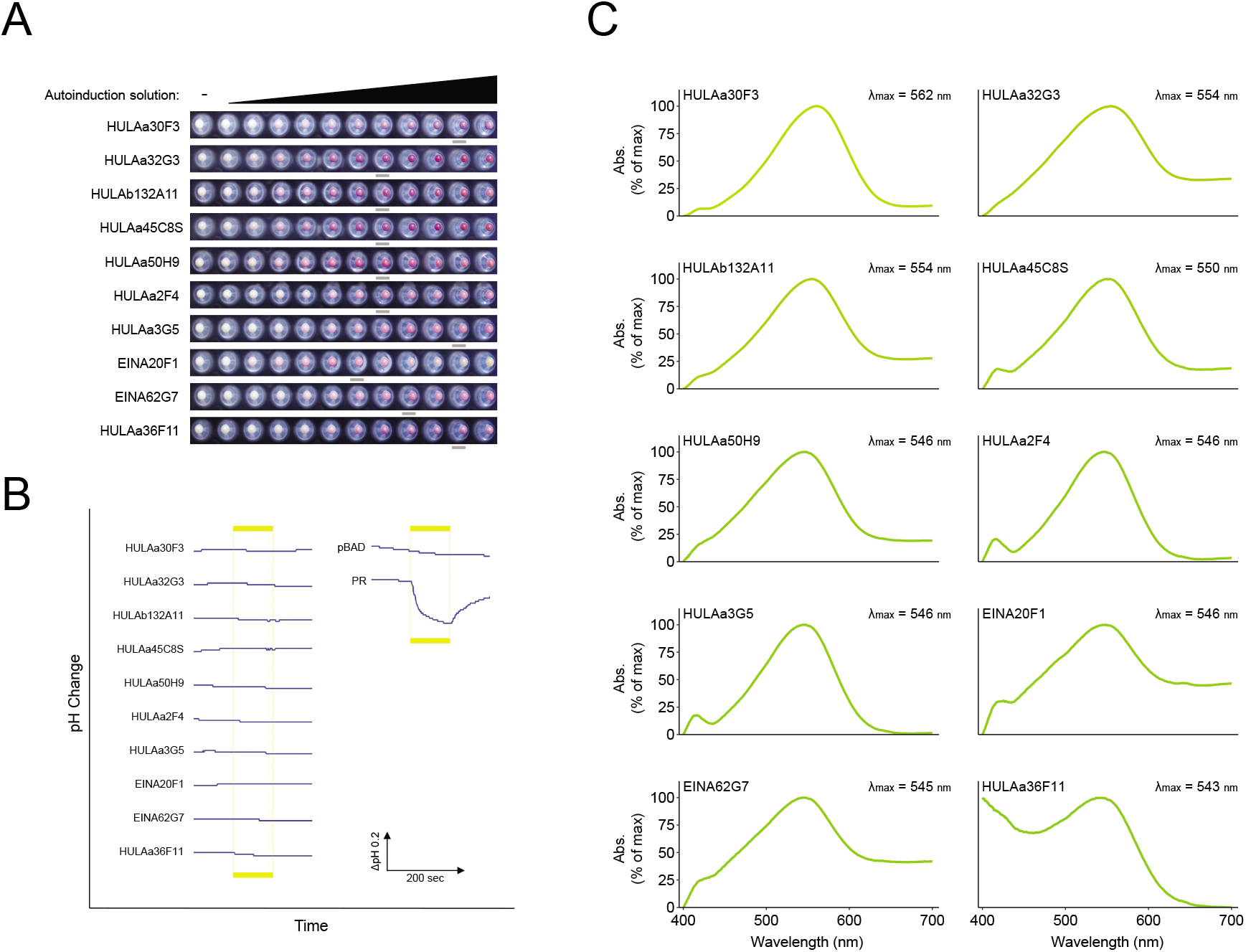
Ion transport activity assay and absorption spectra for HeRs reported in this work. **A**. Optimization of HeRs expression. HeR genes were cloned into pBAD vector and expressed in DH10B™ (Thermo Scientific) cells under different concentrations of autoinduction solution (0-4 µl/ml). Grey bars indicate the optimal expression for each clone. **B**. Monitoring of light-induced pH changes in *E. coli* suspension expressing individual HeRs. Empty vector (pBAD) and a proteorhodopsin pump (clone EINA29G6) served as a negative and a positive control, respectively. Light exposure is indicated by the yellow bars. **C**. Visible absorption spectra of HeRs in DDM-solubilized membranes.

In the first identified HeR-coding environmental clone, 48C12 (Pushkarev et al. 2018), the HeR gene was located at the end of the insert, and was transcribed from the vectors promoter, which hinted at the limited potential of this expression system. In the present screen, HeR genes in eight out of the eleven fosmids were located on the opposite strand or distant from the vector’s promoter (Figure 1A). This finding shows that our screening system is capable of transcribing HeR genes from diverse prokaryotes from their native promoters. Together, the diverse origin of the HeR-coding clones and the expression from their native promoters, highlight metagenomic functional screens as a versatile and powerful tool for characterizing microbial rhodopsins.

Despite the shared biotop, the HeR genes from our screen represented the whole spectrum of HeRs from both of the large phylogenetic subdivisions that are found in diverse prokaryotic taxa (Figure 1 and *SI Appendix*, Fig. S5). HeRs from the monophyletic group A (subfamily 1 in (Kovalev et al. 2020)) are represented mostly by proteins from Actinobacteria, including HeR-48C12 and several of the clones reported here, while group B is more diverse and includes *Bc*HeR (Chloroflexi) and *Ta*HeR (Archaea), even though we could not confirm its monophyletic status in the current study. The two phylogenetic groups demonstrate a tendency to diverge at the loop regions that might reflect divergent functions (see *SI Appendix*, Fig. S5). Nevertheless, cases of horizontal gene acquisitions and replacements even of HeRs from the two divergent groups by closely related taxa are frequent, as exemplified by the fosmid clones from the putative members of the abundant freshwater Actinobacteria of the genus *Planktophila* (see *SI Appendix*, Fig. S2).

Sequence divergence in the HeRs discovered in the current screen provided us with an opportunity to investigate the range of absorption spectra in HeRs in the habitat that we hypothesized would favor spectrally-shifted rhodopsins. Absorption maxima of natural HeR variants characterized thus far have been restricted to the wavelengths between 519 nm and 556 nm. As anticipated for the chosen sampling environment, HULAa30F3 from Lake Ha’Hula, with an absorption peak at 562 nm in DDM-solubilized membranes (Figure 2C) became the most red-shifted natural HeR characterized so far. Based on the ratio between the absorbance of bleached HeR HULAa30F3 and retinal oxime produced (*SI Appendix*, Fig. S6), and molecular extinction coefficient of retinal oxime (ε = 33,600 M-1·cm-1) (Scharf et al. 1992), the ε of HeR HULAa30F3 was estimated to be 46,300 M·cm-1 which is similar to that of other rhodopsins. Somewhat unusual for branch-A HeRs, this clone originated from a Chloroflexi bacterium (Dehalococcoidia) (*SI Appendix*, Fig. S4) and had two of the highly conserved histidine residues (H25 and H81) replaced by the hydrophobic leucine (*SI Appendix*, Fig. S5). Mutations at these positions were previously shown to alter the photocycle of HeR (Pushkarev et al. 2018, Shihoya et al. 2019).

To compare the photocycle of HeR HULAa30F3 with that of typical HeRs (HeR-48C12 and *T*aHeR), we carried out laser flash photolysis measurements (*SI Appendix*, Fig. S7). While a large accumulation of the long-lived O-intermediate was observed from 100-μs to 10-s time region after photo-excitation (Figure 3A and *SI Appendix*, Fig. S7A), the accumulation of the M-intermediate was lower compared to HeR 48C12 and *T*aHeR (Pushkarev et al. 2018, Shihoya et al. 2019). A similar photocycle with a decreased accumulation of the M-intermediate was also observed for the double mutant of the conserved histidines in *T*aHeR (H23F/H82F) (Shihoya et al. 2019). We thus conclude that the two replaced histidine residues are likely responsible for the divergence of the photocycle of HeR HULAa30F3 from that of the typical prokaryotic HeRs and for the lowered accumulation of the M-intermediate, which might be relevant for its molecular function.

**Figure 3.**
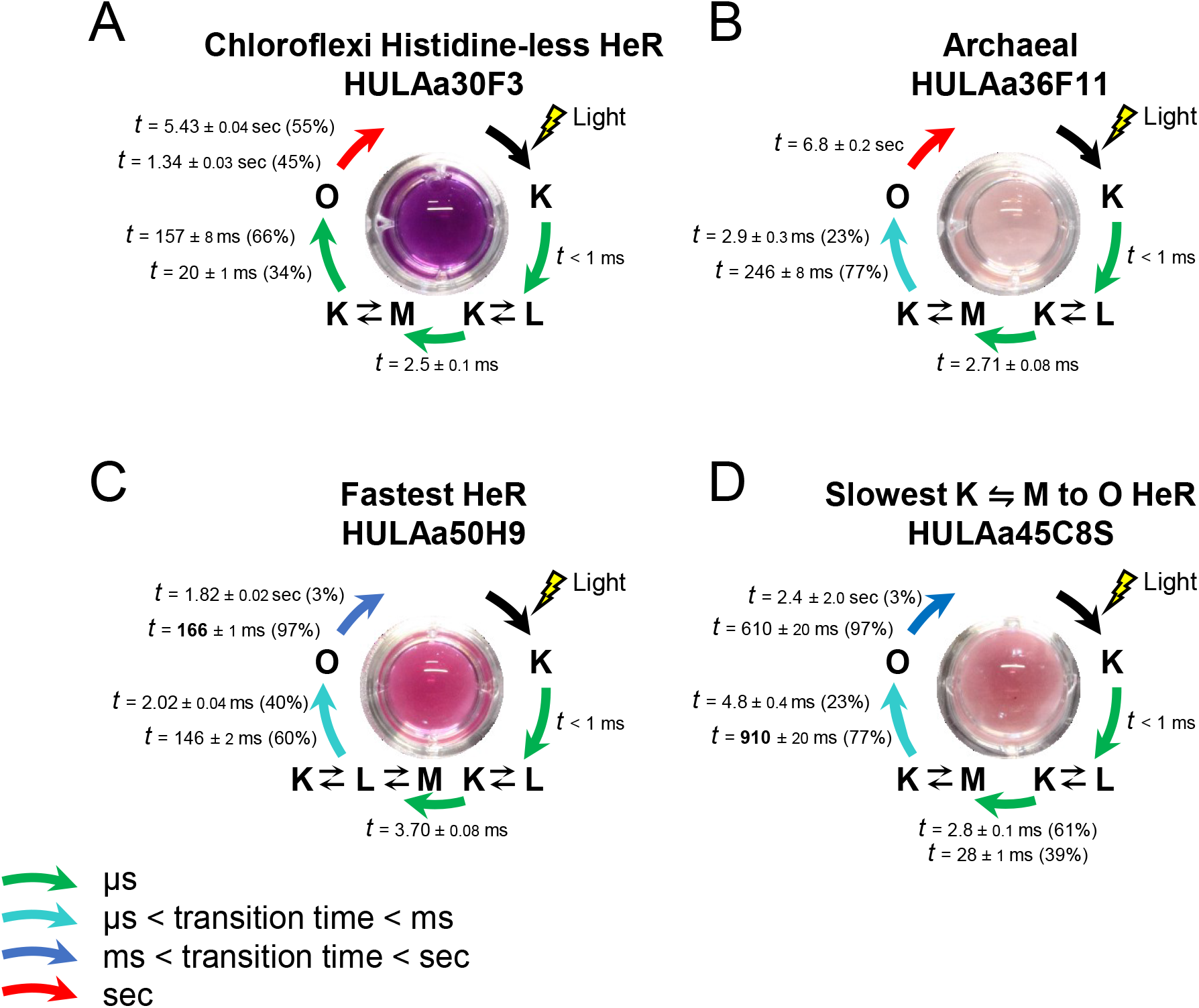
Photocycles of representative HeRs found in this study based on the multi-exponential fitting of the transient absorption changes. The colors of the HeRs purified in DDM are depicted in the centers of each panel. The color of the arrows describes the speed of transition between intermediates according to the legend at the bottom left. HeR HULAa30F3 showed low accumulation of the M-intermediate (see text). While the recovery rate of the initial state of HeR HULAa50H9 from its O is the fastest compared to other HeRs known so far, a large accumulation of the M was observed for HeR HULAa45C8S (*SI Appendix*, Fig. S7D).

The photocycles were also investigated for HeRs HULAa50H9, HULAa36F11, and HULAs45C8S (Figures 3B-D and *SI Appendix*, Fig. S7). Of note, HeR HULAa50H9 showed significantly faster photocycle compared with all known HeRs (97% of protein returned to the initial state with a lifetime *τ* = 166 ms; Figure 3C). At the same time, HeR HULAa45C8S demonstrated the slowest K ⇋ M to O conversion (Figure 3D) (double-exponential process with *τ* = 910 μs and 4.8 ms) that leads to an increased accumulation of the M-intermediate compared to other HeRs.

In the current study we did not attempt to address the question of the molecular function of HeRs. The lack of experimental evidence from previous studies is an indication that the function of the HeRs is not typical for microbial rhodopsins. Nevertheless, to provide a direction for future experiments, we attempted to supplement previously formulated hypotheses about the HeR function(s) with a more solid statistical approach. We observed that environmental fosmid clones provide an immediate genetic context for the target protein genes and this served us as an inspiration for a large-scale search for potential functional associations of HeRs. Together with a dozen of environmental clones, we analyzed a large collection of thousands of HeR-coding DNA segments from genomic and metagenomic assemblies originating from Archaea and several groups of Bacteria known to possess HeR genes, and counted the incidence of protein families, compensating for redundancy (*SI Appendix*, Figs. S8-S9). We reasoned that potential associations between a protein family and rhodopsins can be diverse and not necessarily indicative of a shared molecular function. To highlight the systematic commonalities between HeR neighbors and their differences from type-1 rhodopsins, we designed our analysis around the division of prokaryotic HeRs into the two major phylogenetic groups and their appearance in several unrelated taxonomic groups of prokaryotes (see Figure 1). The HeRs were contrasted against type-1 microbial rhodopsin with a well-predictable function from the same taxonomic groups, i.e. having similar genetic and physiological backgrounds. As this second group of microbial rhodopsins, we chose proteorhodopsins, xanthorhodopsins and related smaller families that are widespread among the three major prokaryotic groups that contributed the bulk of HeR genes for our analysis (*SI Appendix*, Fig. S9). Thanks to the high number of characterized members (ca. 50 in total, six of them from Actinobacteria and Archaea) and a well-studied molecular mechanism, the function of the vast majority of proteins from these families, namely those that have the conserved DxxxTxxxxxxE motif in TM helix C, can be confidently predicted as proton pumps (Beja and Lanyi 2014).

The most common leitmotifs among the protein families encountered in the vicinity of HeR genes in multiple taxa/groups were: (*i*) families with known or plausible connection to oxidative stress, such as SOUL heme-binding domain (e.g. next to HeR HULAa36F11, see Figure 1A), AhpC/TSA family of antioxidants, DNA photolyase, pyridoxamine 5’-phosphate oxidase, tocopherol cyclase; (*ii*) membrane transport proteins: ABC and MFS transporters; (*iii*) proteins involved in translation: SpoU rRNA methylase, various tRNA-modifying enzymes, S30EA ribosomal protein and (*iv*) diverse protein families that utilize light for their activity: DNA photolyase, cobalamin-binding proteins and PAS-domain-containing proteins (Figure 4 and *SI Appendix*, Fig. S10, Dataset S3). Nevertheless, none of these families appeared unique to HeRs as all of them could be found associated with the DTE proton pump genes as well. In fact, association of photolyases with some DTE proton pumps in Actinobacteria has been previously reported as well (Ghai et al. 2013).

**Figure 4.**
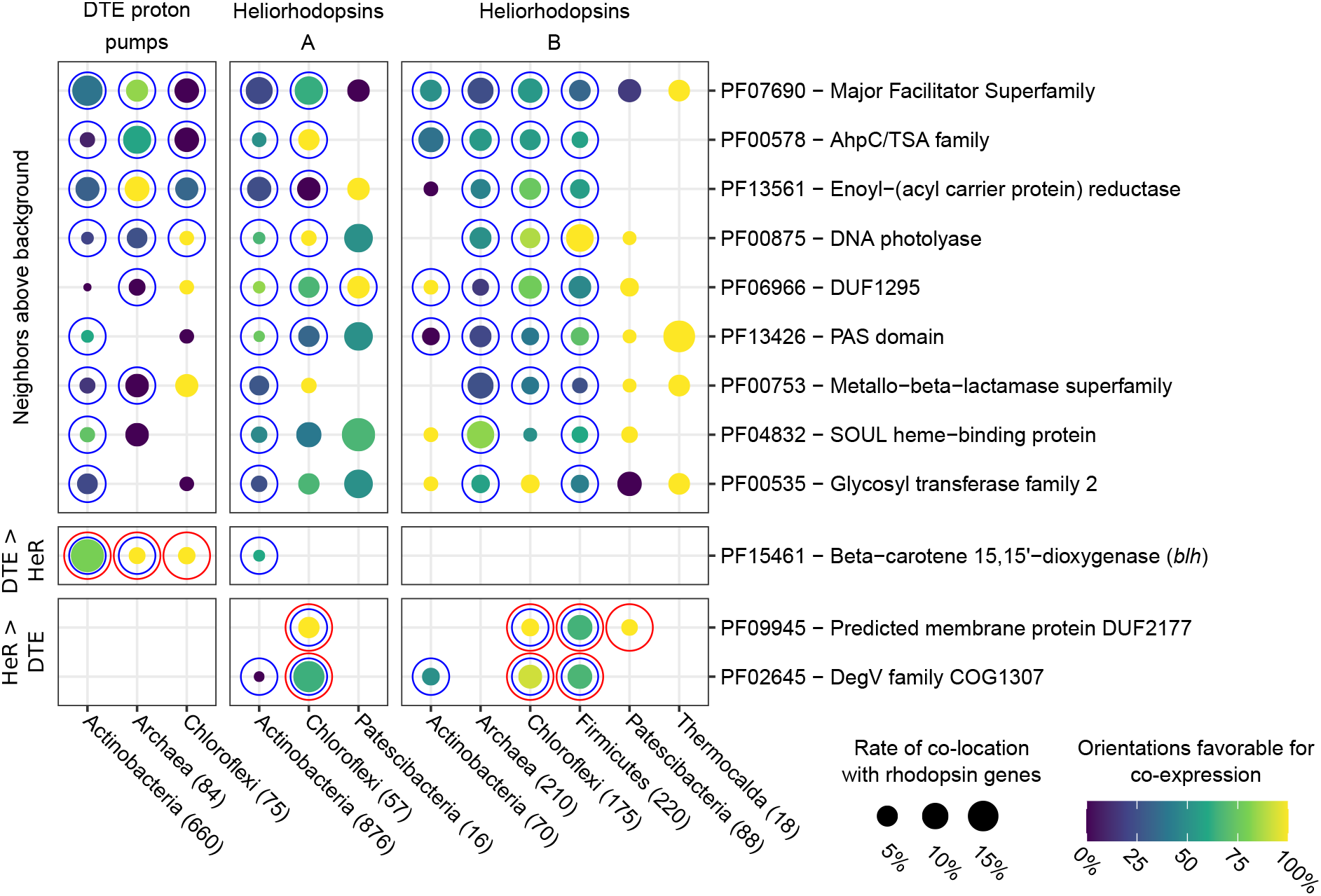
Analysis of protein families in genes neighboring prokaryotic HeRs and putative rhodopsin proton pumps. Prokaryotic HeRs are subdivided into the two phylogenetic groups, A and B (see Figure 2A and *SI Appendix*, Fig. S5), and major taxa with numbers in parentheses indicating the number of non-redundant HeRs analyzed. The matches are grouped by the pattern of their appearance: 1) genes commonly appearing in the vicinity of HeRs and DTE proton pumps with rates above background but without preference for either rhodopsin type; 2) genes disproportionately present in the vicinity of proton pump genes; 3) genes disproportionately present close to HeR genes. The size of the dots corresponds to the proportion of distinct rhodopsins with the corresponding neighbors, the color – to the incidence of gene orientations indicative of potential co-expression or co-regulation. Blue circles indicate significant over-representation over background, while red circles indicate significant differences between the rhodopsin types. The outline of the pipeline is presented in *SI Appendix*, Fig. S8. The exact filtering applied for this figure is described in Materials & Methods. The full listing of the protein families and corresponding tests can be found in Dataset S3.

One of the surprising discoveries was the rarity of the *blh* (β-carotene 15,15’-dioxygenase) genes required for the final step of the retinal biosynthetic pathway in prokaryotes among HeR neighbors. Despite a rather moderate presence of *blh* among the neighbors of DTE proton pumps in Actinobacteria, Chloroflexi and Archaea, it still appeared as their single most characteristic neighbor when compared to HeR neighbors (see Figures 4 and *SI Appendix*, Fig. S10). To confirm that this pattern is generalizable to whole genomes, and not only to the surroundings of HeRs, we correlated the presence of HeR genes and *blh* in complete uncontaminated assemblies (*SI Appendix*, Fig. S11) to come to the conclusion that HeR co-occur with *blh* mostly in cases when the same genomes already encode other microbial rhodopsins. This provides support to the idea of the existence of an as yet unknown pathway for retinal biosynthesis as has been recently demonstrated for the freshwater actinobacterium *Aurantimicrobium minutum* KNC (Nakajima et al. 2020). Notice that this bacterium harbors not only a DTE proton pump (xanthorhodopsin) but also a HeR.

The list of the protein families that systematically differentiate between HeRs and DTE proton pump neighbors was in fact very short. The two protein families that had a significant presence (>5%) in at least two groupings of HeRs, and demonstrated a high rate of co-expression potential (>50%) in at least three groupings were: membrane protein of unknown function DUF2177 and fatty acid-binding protein DegV (see Figure 4) (a DegV-family gene was found e.g. next to HeR HULAa30F3, see Figure 1A). Most of the contribution to this pattern was provided by HeRs from Firmicutes and Chloroflexi, and to a lesser extent Actinobacteria and Patescibacteria. Interestingly, the presence of the two protein families in the vicinity of HeR genes not only crosses taxonomic borders, but also unites both of the phylogenetic groups of prokaryotic HeRs (see above), signifying thus a very systematic appearance. In fact, DUF2177 and DegV often co-appear in the vicinity of the same HeR genes accompanied by a third gene, from the DUF1295 family that alone has a sporadic appearance among DTE rhodopsin neighbors as well (see Figure 4). Summary statistics for the genome-wide co-occurrences confirms this pattern: while DUF1295 and DegV often co-occur with other microbial rhodopsins in the same genome, the particular combination of the three protein families DUF2177, DegV and DUF1295 is characteristic to HeR-coding genomes (18% of HeRs appear in genomes with all three protein families) and is virtually absent from the genomes harboring other microbial rhodopsins in this prokaryotic sample. Among the three protein families, DegV has the highest co-appearance with HeR (59%) genome-wise (see *SI Appendix*, Fig. S11).

Proteins from the DegV family are soluble and characterized members (FakBs) participate in activation of unphosphorylated fatty acids (FA) by binding them with high affinity and exposing to a specific kinase (FakA, from the family of dihydroxyacetone kinase Dak2) for phosphorylation (Parsons et al. 2014, Broussard et al. 2016). The ability of DegV proteins to bind FAs is universal across the family and is determined by a conserved pocket, the shape of which defines selectivity for different FAs (Broussard et al. 2016). Dak2-DegV fusion proteins have been reported (Erni et al. 2006), and such fusions are occasionally encountered also in the vicinity of HeR genes in Chloroflexi and Firmicutes as well (see *SI Appendix*, Fig. S10). In this context, it is interesting to note that genes coding for enoyl-acyl carrier protein reductase, an enzyme involved in type-II FA synthesis, were also encountered in the vicinity of HeR genes in multiple groupings, but at the same time had a significant presence among neighbors of DTE proton pumps as well (see Fig. 4). The second family characteristic of HeR neighbors, DUF2177, includes membrane proteins of unknown function and was also entirely absent among neighbors of DTE rhodopsin pumps.

The functional relationship between these two protein families and HeRs is unclear especially since even this association is not universal among HeR genes. Notice however that reminiscently of the DegV proteins, the crystal structures of HeRs revealed that they bind such lipids from the membranes used for crystallization as the FA monooleate and the alkane eicosane and that the corresponding binding site is localized to the fenestration that exposes the β-ionone ring of the retinal moiety (Shihoya et al. 2019, Kovalev et al. 2020). Taken together, these observations are suggestive of HeRs participating in FA modification in a light-dependent manner. This idea, in turn, might provide a logical connection to the fact that HeRs are not encountered in classical diderm bacteria. Sensitivity to photosensitizers is one of the most pronounced physiological differences between monoderms and diderms: the lipopolysaccharide-containing outer membrane serves as a barrier to such photosensitizers as natural porphyrins that otherwise under illumination cause production of reactive oxygen species (ROS) in the cellular membrane (Malik et al. 1990, Sperandio et al. 2013). It is thus possible that the co-location of HeR genes with the genes implicated in oxidative stress is explained not only by a common regulation by light exposure as must be assumed for the proton pumps (see Ghai et al. (2013)). We therefore hypothesize that prokaryotic HeRs might have a light-dependent enzymatic activity related to repairing oxidative damage in the cellular membrane or to membrane remodelling aimed at preventing such damage.

It has been almost three years since the discovery of HeRs, yet their function remains elusive. Our search for new and expressible HeRs combined with biophysical characterization, phylogenomics and co-expression potential analyses give further leads to the function of this diverse and widespread family.

## Materials and Methods

### DNA sampling for library preparation

Freshwater sampling from Ein Afek reserve (Israel) was performed on 25/11/2019 (32°50’44.15”N 35° 6’49.04”E). Twenty liters of water from the surface were filtered through GF/D filters (Whatman) and collected on a 0.22 µm Durapore filter (Millipore). The Durapore filter was suspended in a lysis buffer (50 mM Tris-HCl, pH 8.0, 40 mM EDTA, pH 8.0, 0.75 M sucrose) and flash frozen using liquid nitrogen on site.

Peat lake sampling was performed on 18/10/2018 in Agamon Ha’Hula peat lake, Israel at two locations: A – the main water body (33°06’22.4”N 35°36’09.4”E), and B – a shallow pond (33°06’24.4”N 35°36’04.7”E). Forty liters of water from 20 cm depth were filtered through GF/D and a 0.22 µm Durapore filter (Millipore). DNA from both sample sites was extracted from the filter using a phenol-chloroform protocol (Wright et al. 2009).

Fosmid libraries of peat lake and freshwater samples were constructed with a pCC2Fos copy control library kit according to manufacturer’s protocol (Epicentre Biotechnologies, Cat. No. CCFOS059), and 198 96-well plates (66 from each sampling location) were stored in LB-glycerol 7% at −80 °C.

### Rhodopsin screen

Cells were inoculated from fully thawed library plates into 96-well 2.2-ml plates (ABgene, Cat. No. AB-0932) filled with 1 ml LB supplemented with 25 μg/ml chloramphenicol and 2 µl/ml CopyControl fosmid autoinduction solution (Epicentre Biotechnologies, Cat. No. AIS107F). The plates were grown at 30 °C shaking at 700 RPM for 16 h covered by an AeraSeal gas-permeable sheet (EXCEL Scientific Cat. No. BS-25) after which all*-trans* retinal (Sigma Cat. No. R2500) was added to a final concentration of 50 μM and the plates were incubated in the dark at RT with shaking at 100 RPM for 4 h. Cells were pelleted by centrifugation at 3000 RCF for 10 min, washed twice with 500 μl salt mix (10 mM NaCl, 10 mM MgSO_4_ and 100 μM CaCl_2_) and collected for visual color evaluation.

Clones showing color phenotype were re-screened in the presence/absence of retinal to show retinal dependent pigment manifestation. Each clone was inoculated into four wells in a flat-bottom 96-well plate (Thermo Scientific Cat. No. 167008) containing 7% LB-glycerol supplemented with 25 μg/ml chloramphenicol and 10 μg/ml streptomycin and incubated overnight at 37°C after which stored at -80°C. Cells were inoculated from a fully thawed re-screen plate into 96-well 2.2-ml plates and grown as described. Retinal was added to two out of four duplicates and after incubation (same as with the first screen) and centrifugation to pellets, washed once with 500 μl salt mix (10 mM NaCl, 10 mM MgSO_4_ and 100 μM CaCl_2_) and transferred to 96 well transparent PCR plates (Axygen, Cat. No. PCR-96-FLT-C) for fixation with 1% agarose.

### Sequencing and *de-novo* assembly

Library preparation was done using the Nextera XT sample prep kit (Illumina) according to manufacturer’s protocol. The libraries were sequenced with HiSeq 2500 and MiSeq (Illumina) sequencers to obtain 150-nt paired-end reads. After optional quality and vector trimming, the reads were assembled using SPAdes v. 3.5.0 (Nurk et al. 2013). ORF prediction and functional annotation were performed with PROKKA v. 1.14.6 (Seemann 2014) in the metagenome mode with domain-specific settings and applying a cutoff of 10^−5^ on E-value and 80% on coverage for comparisons with references. Fragmented ORFs at the ends were annotated with the aid of MetaGeneMark (Zhu et al. 2010). Fosmid assemblies were submitted to GenBank under the accession numbers MW122873-MW122884.

### Expression optimization

Unique HeRs were cloned into pBAD vector with an N-terminal 6xHIS tag. Fresh colony was inoculated into 96-well 2.2-ml plates (ABgene, Cat. No. AB-0932) filled with 1 ml LB supplemented with 25 μg/ml chloramphenicol and CopyControl fosmid autoinduction solution at concentrations ranging from 4 to 0.004 µl/ml and shaken overnight at 30°C with 750 RPM. All-*trans* retinal (50 µM final) was added, and were incubated in the dark at RT with shaking at 100 RPM for 4 h. Cells were collected by centrifugation and washed twice with salt solution (10 mM NaCl, 10 mM MgSO_4_ and 100 µM CaCl_2_). Optimal expression was determined visually based on uniformity and intensity of color.

### Absorption spectra analysis

Expression-optimized *E. coli* cultures (50 ml) were washed twice with salt solution (10 mM NaCl, 10 mM MgSO_4_ and 100 µM CaCl_2_). Cells were resuspended in a 25 ml salt solution containing protease inhibitor cocktail 1 µl/ml (Sigma-Aldrich Cat. No. P8849) and PMSF 1 µM (Sigma-Aldrich Cat. No. P7626) and disrupted by 10 passes in a microfluidizer at 60 psi. Cells were centrifuged at 4°C at 5000 RCF for 10 min to pellet cell debris and the supernatant was centrifuged at 4°C at 37,000 RCF for 1 h to pellet the membranes. Membranes were solubilized overnight in a buffer containing 50 mM MES, 300 mM NaCl, 5 mM imidazole, 5 mM MgCl_2_, 2% DDM, pH 6.5. DDM solubilized proteins were centrifuged at 4°C at 20,000 RCF for 10 min and the supernatants were taken for spectral characterization. Absorption spectra were measured with a Shimadzu UV-1800 spectrophotometer.

### Laser flash photolysis of HeRs

The transient absorption change after the photo-excitation of HeR was investigated by laser-flash photolysis method (Pushkarev et al. 2018). The *E. coli* cells expressing HeR (C43(DE3) strain for HeR HULAa30F3 and HULAa50H9, and DH10B™ (Thermo Scientific) for HeR, HeR HULAa36F11 and HeR HULAa45C8S) were suspended in 100 mM NaCl, 50 mM Tris-HCl (pH 8.5). Then, the cells were treated with 1 mM lysozyme and disrupted by sonication for 10 min, three times. The large membrane fraction and undisrupted cells were removed by centrifugation (22,262 RCF, 10 min). The optical density of the suspension was adjusted to be 0.8-0.9 by dilution. The sample solution was illuminated with a second harmonics generation of a nano-second pulsed Nd^3+^-YAG laser (*λ* = 532 nm, INDI40, Spectra-Physics) with the pulse energy of 5.1 mJ/cm^2^ pulse. The transient absorption spectrum of rhodopsin after the laser excitation was obtained by measuring the intensity of white light that passed through the sample before and after laser excitation at *λ* = 350-750 nm with an ICCD linear array detector (C8808-01, Hamamatsu). Whereas the detector and the excitation laser were electronically synchronized in the microsecond time region, the jitter between laser pulse illumination and detector exposure could not be so precisely controlled due to instrumental limitation during the scanning of the time-region of *t* > 70 ms and *t* > 1 s for HULAa36F11 and HULAa45C8S, respectively, showing longer photocycles. In the latter case, we needed to place a notch filter in front of the detector to avoid the saturation by scattered laser pulse, and thus the absorption change could not be observed at *λ* = 523-540 nm. To increase the signal-to-noise ratio, 60 signals were averaged and singular value decomposition analysis was applied (Chizhov et al. 1996). The time evolution of transient absorption change at specific wavelengths after photo-excitation was measured by monitoring the change in intensity of the monochromated output of an Xe arc lamp (L9289-01, Hamamatsu Photonics) passed through the sample suspension by a photomultiplier tube (R10699, Hamamatsu Photonics) equipped with a notch filter (532 nm, bandwidth = 17 nm, Semrock) to remove scattered pump pulse. The signals were monitored and stored by a digital oscilloscope (DPO7104, Tektronix). We averaged 30-60 signals.

### Classification of environmental clones

Rough phylogenetic position of the fragments was first assessed by performing searches against translated nucleotide sequences of reference assemblies included in GTDB r. 95 (Parks et al. 2018) using fosmid protein sequences as queries with ublast (E-value threshold 1e-10) from usearch v11.0.667 (Edgar 2010). For each reference assembly-fosmid pair counts of homologous genes and the corresponding average identities were obtained and phylogenetic affinities of the most similar assemblies were assessed. At this step, the clone HULAa45C8S (Actinobacteria, putative Micrococcaceae) was excluded from further analysis as too short and clones HULAa55C9 and HULAa32G3 (Actinobacteria) were rejected because of low average identity values and high diversity of the best-matching reference assemblies. For each of the chosen taxonomic clades, orthologous groups suitable for phylogenetic analysis were obtained using proteinortho v. 6.0.25 (Lechner et al. 2011) with ublast as the search program, requiring a minimum of 60% identity for pairwise matches and a threshold of 1.0 on the minimum algebraic connectivity. Orthogroups with paralogues or less than five members were filtered out. For each orthogroup, phylogeny was reconstructed by aligning the protein sequences with mafft v. 7.453 (localpair mode), trimming the alignments with trimAl (automated1 mode) and obtaining gene trees with iqtree v. 1.6.12. To increase robustness of the subsequent species tree inference, gene trees were also obtained for the reference assemblies from the 120 or 122 marker gene alignments included in GTDB. The gene trees were further filtered with treeshrink v. 1.3.6 (Mai and Mirarab 2018) and the species trees were inferred with ASTRAL v. 5.7.4 (Zhang et al. 2018). Branch lengths were estimated by ERaBLE v. 1.0 (Binet et al. 2016) and the root, taxonomic assignments and support values were transferred from the GTDB reference tree. To enable cross-comparisons, the metagenomic contigs containing previously characterized HeRs 48C12 and *Ta*HeR were included in this analysis as well. The pipeline is available via https://github.com/BejaLab/FosmidPlacer.

### Phylogenetic analysis and classification of HeRs

A detailed outline of the approach taken for phylogenetic reconstruction and classification of prokaryotic HeRs is presented in *SI Appendix*, Fig. S8. Sequences for the phylogenetic analysis of HeRs were collected from genomic and metagenomic assemblies (see below) and Uniprot. Initial experiments confirmed the disproportionately high rates of evolution in the putative superclade of viral and eukaryotic HeRs (Kovalev et al. 2020), and they were removed from the analysis, focusing thus on the prokaryotic HeRs only. Sequences were clustered in two steps: at 80% and subsequently at 60%, with CD-HIT v. 4.8.1 (Li and Godzik 2006), aligned with mafft (localpair) and trimmed with trimAl to filter out positions occupied by gaps in more than 10% of the sequences. Maximum likelihood phylogeny was reconstructed using iqtree (Minh et al. 2020) using default settings. To guarantee convergence, the search was repeated 20 times with different seeds and the highest-likelihood tree was selected. LG+F+R7 was selected as the best-fit substitution model. Branch supports were obtained by ultrafast bootstrap with 1000 replicates.

### Analysis of gene neighbors

The material for the analysis of rhodopsin neighbors was obtained from all available NCBI wgs projects, assemblies and environmental clones in GenBank (Dec 2020) assigned to one of the major prokaryotic groups known to harbor HeRs: Archaea, Actinobacteria, Chloroflexi, Firmicutes (incl. Tenericutes), selected taxa from Patescibacteria (= Candidate phyla radiation) and three phyla from the Thermocalda clade (Dictyoglomi, Thermotogae and Caldisericota (Cavalier-Smith and Chao 2020)) (see *SI Appendix*, Fig. S9 for details). For the sake of exhaustiveness, the initial search for HeR-containing genomic fragments was performed by fetching translations of all ORFs and matching them against the Pfam profile of HeR (PF18761.2) with hmmsearch from HMMER v. 3.3.2 (Eddy 1998) (see *SI Appendix*, Fig. S8A). An analogous search was performed for putative proton pumps with the conserved motif DTE using a protein profile prepared from UniRef90 sequences matching GPR (Q9F7P4) and/or XR (Q2S2F8) (available in Dataset S4). To avoid spurious matches and pseudogenes, HeR hits were required to have two most conservative positions intact: a lysine in TM helix 7 and an arginine in TM helix 3. The putative DTE proton pumps were required to have a lysine in TM helix 7 and the DTE motif in TM helix 3. Genes were annotated for the matching DNA segments with GeneMarkS v. 4.32 (Besemer 2001) if not provided in the source database and the rhodopsin genes were re-annotated using hmmsearch followed by manual curation when needed. Genes within windows of ±10 in both directions around the rhodopsin genes were searched against Pfam r. 33.1 with pfam_scan.pl (Finn et al. 2014). In order to be considered non-redundant, a rhodopsin gene with its neighbors was required to have a unique ordered set of three 80%-identity clusters for the rhodopsin itself and its two immediate neighbors. For the non-redundant rhodopsin genes, for each gene with a Pfam match the pipeline collected the distance to the rhodopsin gene and whether the matching gene was potentially co-expressed/co-regulated with the rhodopsin gene (*SI Appendix*, Fig. S8B). In addition, for rhodopsins coming from long (meta)genomic assemblies (≥ 1000 genes), the incidence of the Pfam families was assessed genome-wide. The collected Pfam family matches were summarized per taxon per rhodopsin group (HeR-A, HeR-B and DTE proton pumps). Matching families were pre-filtered to include only those that appear in >5% of the windows, distanced on average less than four genes away from the corresponding rhodopsin gene and potentially co-express/co-regulate with the rhodopsin in >10% of the cases, each requirement to be fulfilled in at least one grouping. For each Pfam family its incidence among HeR neighbors was compared against that for the proton pump neighbors (see *SI Appendix*, Fig. S8C for details). Comparisons were also performed against the background for each rhodopsin grouping by contrasting the number of the genes matching a particular Pfam family in the vicinity of the rhodopsin genes against the counts for the rest of the genome (see *SI Appendix*, Fig. S8C). The counts were compared using Fisher’s exact test with p-value correction according to Holm’s method.

Genome-wide incidence of gene co-occurrences of HeRs and other rhodopsins with a chosen set of protein families was estimated for representative genome assemblies included in GTDB. A simple rule “difference between CheckM completeness and contamination of at least 95” was applied to select high-quality genomes. The genomes were annotated with GeneMarkS if no annotations were available in NCBI Assembly, and searched for rhodopsin genes and other selected protein families with pfam_scan.pl. A simplified approach to detecting rhodopsin genes was applied here, with matches against PF18761 (HeR) or PF01036 (other microbial rhodopsins) Pfam profiles required to pass the recommended thresholds. Note that the PF01036 profile effectively matches a wide range of rhodopsin families encountered in prokaryotes but is not sensitive enough to detect all of them.

For the figure representation (Figure 4), the following customized filtering of the Pfam protein families was applied to highlight hits with the most prominent presence among rhodopsin neighbors. For HeRs, the protein families were required to be significantly more abundant among HeR neighbors with respect to DTE proton pump neighbors (adjusted *p*-values < 0.01) in at least three out of nine groupings (combinations of taxa and HeR phylogenetic group), have an orientation favorable for co-expression with HeR genes in > 50% of the cases in ≥ 3/9 of the groupings, be present in the vicinity of > 5% of the HeR genes in ≥ 2/9 of the groupings, and > 2% in ≥ 4/9 of the groupings. For DTE proton pumps, the following thresholds were set: significantly higher presence with respect to HeR neighbors in at least two out of three taxa, have an orientation favorable for co-expression in > 50% of the cases in ≥ 2/3 of the taxa, be present in the vicinity of >5% of the HeR genes in ≥ 2/3 of the taxa, and >2% in ≥1/3 of the taxa. Protein families that had insignificant differences in their presence between neighbors of HeR and DTE proton pumps but had above-background presence were filtered as follows: significantly higher presence in the vicinity of rhodopsin genes with respect to background in ≥ 4/12 of the rhodopsin groupings (nine HeR groupings and three taxa with DTE proton pumps), orientation favorable for co-expression in >50% of the cases in ≥4/12 groupings, presence in the vicinity of >5% of rhodopsin genes in ≥ 3/12 of the groupings, and >2% in ≥ 5/12 of the groupings. The full list of protein families per grouping is provided in Dataset S3.

## Supporting information

Dataset S1 Fosmids GenBank

Dataset S2 Protein sequences

Dataset S3 Neighbors

Dataset S4 hmm

## Data Availability

Annotated sequences of the fosmids reported here are available from GenBank under accessions MW122873-MW122884 (also provided as Dataset S1). Sequences collected for the analysis of rhodopsin neighbors with corresponding metadata are included as Dataset S2. Summary statistics results of statistical analysis of protein families among rhodopsin neighbors are provided in Dataset S3.

## Code Availability

Code used for the bioinformatic analyses is available from https://github.com/BejaLab/Hula-Heliorhodopsins. The pipeline used for phylogenetic placement of fosmids is available as a separate repository: https://github.com/BejaLab/FosmidPlacer.

## Author contribution

A.P. and O.B. conceived the project, A.C., R.T., S.L., S.Y., O.B., and A.P. collected, constructed and screened the libraries. A.R performed bioinformatic analyses. K.M., T.N., and K.I. performed biophysical analyses. A.C., A.R., K.I., O.B., and A.P. wrote the manuscript.

## Acknowledgments

We thank Idan Barnea for help with sampling in Agamon Ha’Hula and the Keren Kayemeth LeIsrael – Jewish National Fund (KKL-JNF), the acting Forest Service of Israel for the permit to sample in Agamon Ha’Hula, and Israel Nature and Parks Authority for the permit to sample in Ein Afek reserve. This work was supported by Israel Science Foundation – F.I.R.S.T. (Bikura) Individual grant (3592/19 to O.B.), Grants-in-Aid from the Japan Society for the Promotion of Science (JSPS) for Scientific Research (KAKENHI grant Nos. 17H03007 and 20K21383 to K.I.), and the Japan Science and Technology Agency (JST), PRESTO, Japan (grant No. JPMJPR1888 to T.N.). O.B. holds the Louis and Lyra Richmond Chair in Life Sciences.

## Conflict of Interest

The authors declare no conflict of interest.

## Notes

https://github.com/BejaLab/FosmidPlacer

https://github.com/BejaLab/Hula-Heliorhodopsins

### Competing Interest Statement

The authors have declared no competing interest.

