## Supplemental Figures for "Characterization of heliorhodopsins detected via functional metagenomics in freshwater Actinobacteria, Chloroflexi and Archaea"

**This PDF file includes:**

Figures S1 to S11

**Other supporting information includes:**

**Dataset S1 (flat GenBank format)**. Annotated sequences of the reported fosmids and proteorhodopsin gene used as positive control in Figure 2B.

**Dataset S2 (Excel spreadsheet).** Protein sequences and metadata for prokaryotic HeR and DTE proton pump genes collected for the analysis of gene neighbors.

**Dataset S3 (Excel spreadsheet)**: Summary statistics and results of Fisher's exact test for the different Pfam protein families and domains that deviate significantly between groups of rhodopsins (heliorhodopsins vs. DTE proton pumps) or between the vicinities of rhodopsin genes and background genomic locations.

**Dataset S4 (flat hmm file):** Protein profile used to collect DTE proton pumps, built from proteorhodopsin and xanthorhodopsin sequences in UniRef90. Note that the matches were required to have the conserved DTE motif in TM helix C and lysine in TM helix G.

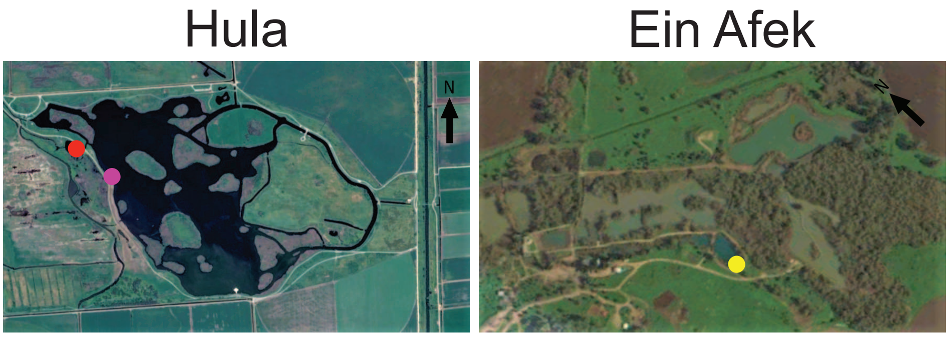


**Figure S1. Sampling sites.** Colored dots represent the different sampling sites: pink – Hula A (33°06'22.4"N 35°36'09.4"E), red – Hula B (33°06'24.4"N 35°36'04.7"E), and yellow – Ein Afek (32°50'44.15"N 35° 6'49.04"E). Figures were taken from Google Earth Pro 7.3.3 (December 20, 2014). North of Israel. Positions as mentioned above, eye altitude 762 m.

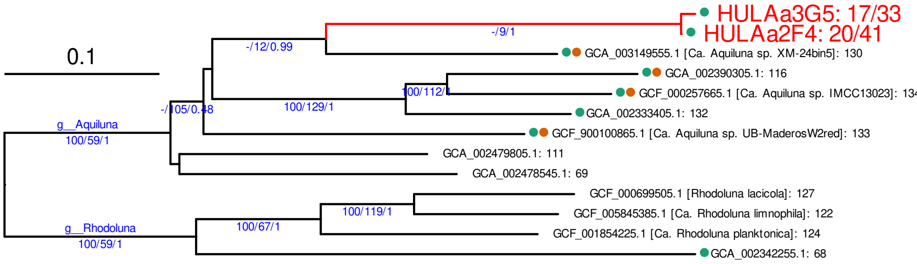


Bacteria, Actinobacteria: the Aquiluna/Rhodoluna clade

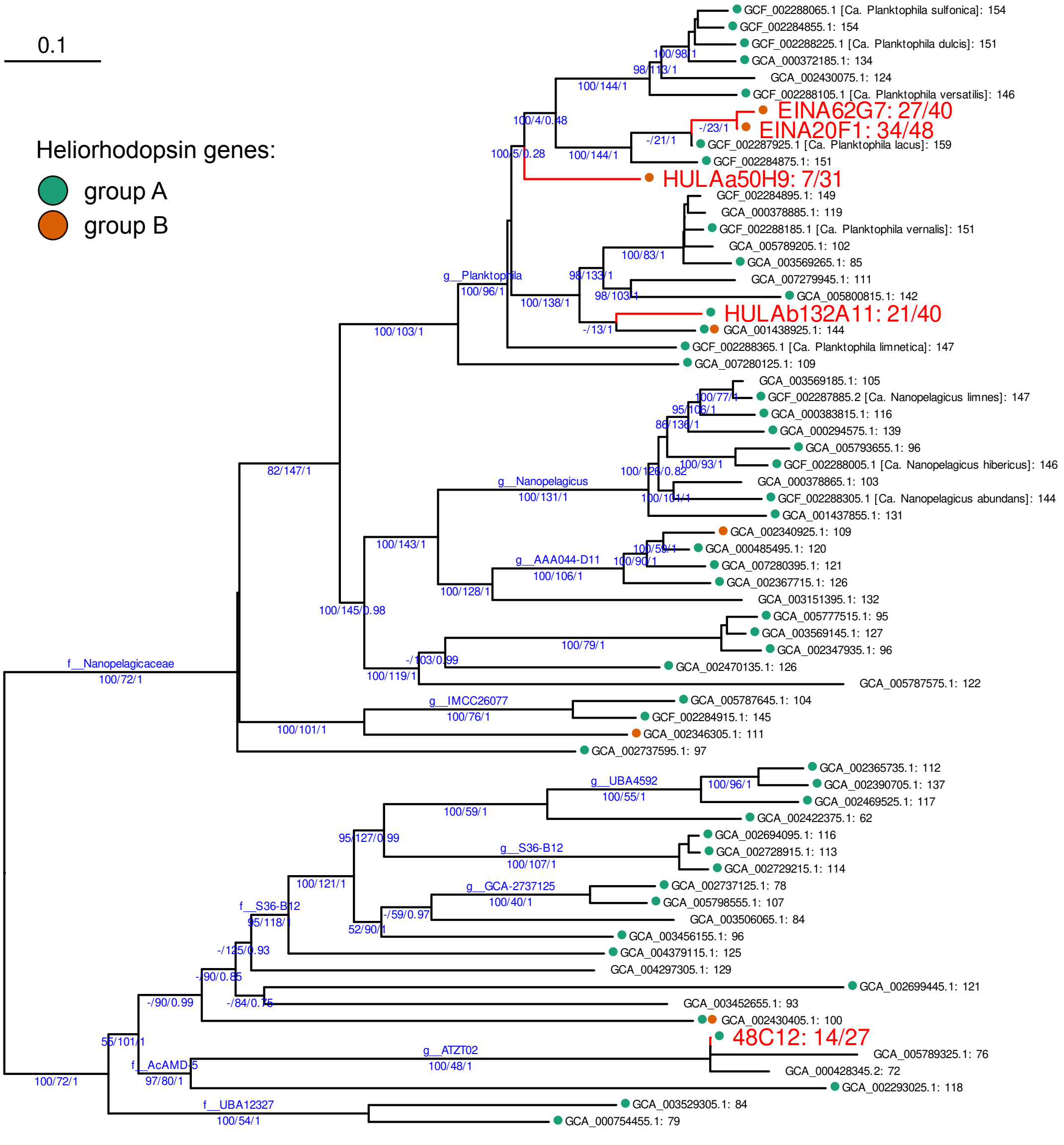


Bacteria, Actinobacteria: Nanopelagicales

**Figure S2**. **Phylogenetic affinities of six of the actinobacterial clones isolated in this study.** Estimated phylogenetic position of the actinobacterial clones including the previously reported clone 48C12. For named branches present in GTDB the corresponding taxa are provided above the branches. Below the branches the three numbers indicate: bootstrap branch support from the GTDB reference tree, effective number of genes in the species inference and local posterior probability (notice that for low values of the effective number of genes, posterior probabilities are not informative). The tips are labelled with NCBI assembly accessions (in black) or clone names (in red). For the environmental clones the two numbers after the colon indicate the number of the genes taken for gene phylogenies (this number could decrease further due to filtering) and the total number of the genes on the contig. Dots indicate the presence of HeR genes from the two phylogenetic groups.

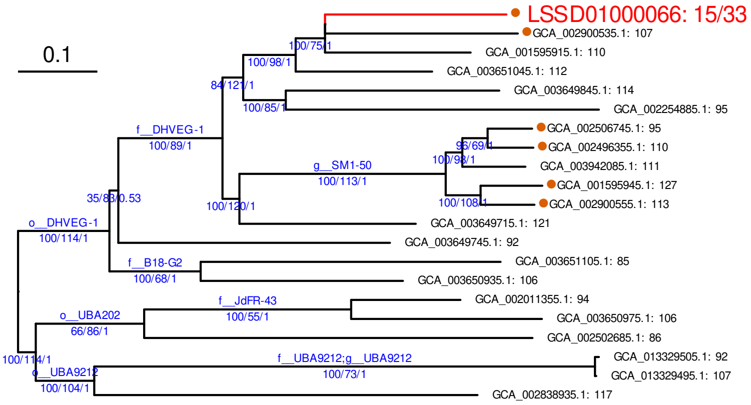


Archaea, Thermoplasmatota: class E2

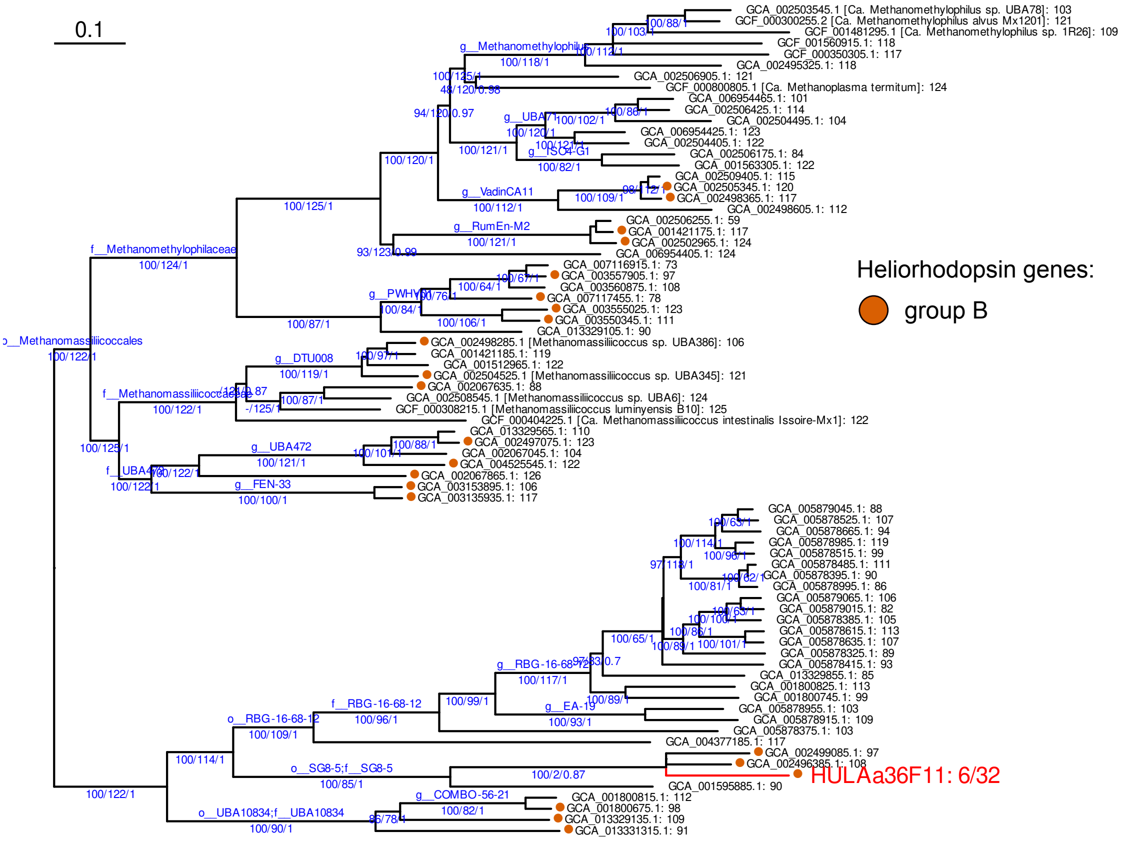


Archaea, Thermoplasmatota: class Thermoplasmata

**Figure S3**. **Phylogenetic affinities of the archaeal clone isolated in this study.** For comparison, the same analysis was performed for the metagenomic contig that contained *Ta*HeR (GenBank accession number LSSD01000066). See Supplementary Figure S2 for further details.

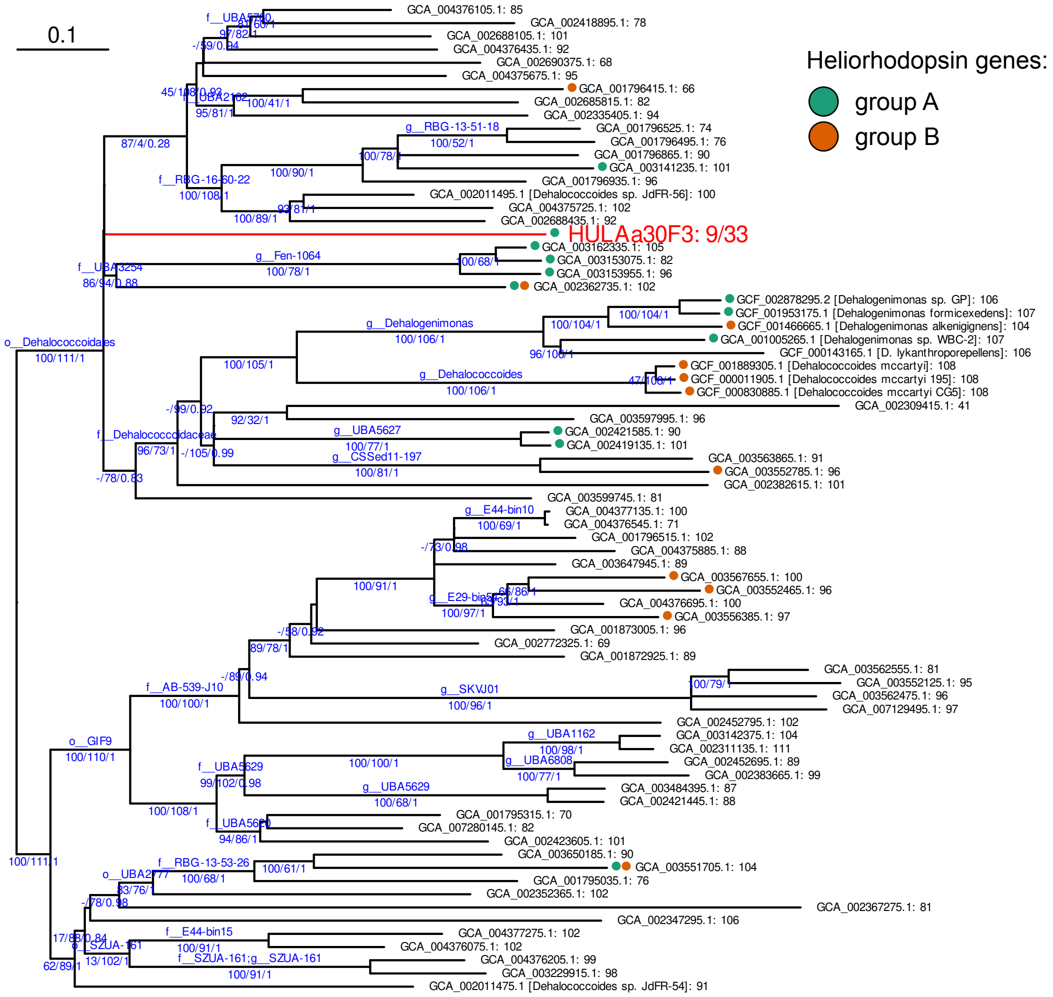


Bacteria, Chlroflexi: the clade uniting the orders Dehalococcoidales, SZUA-161, UBA2777 and GIF9

**Figure S4**. **Phylogenetic affinities of the chloroflexal clone isolated in this study.** See Supplementary Figure S2 for description details.

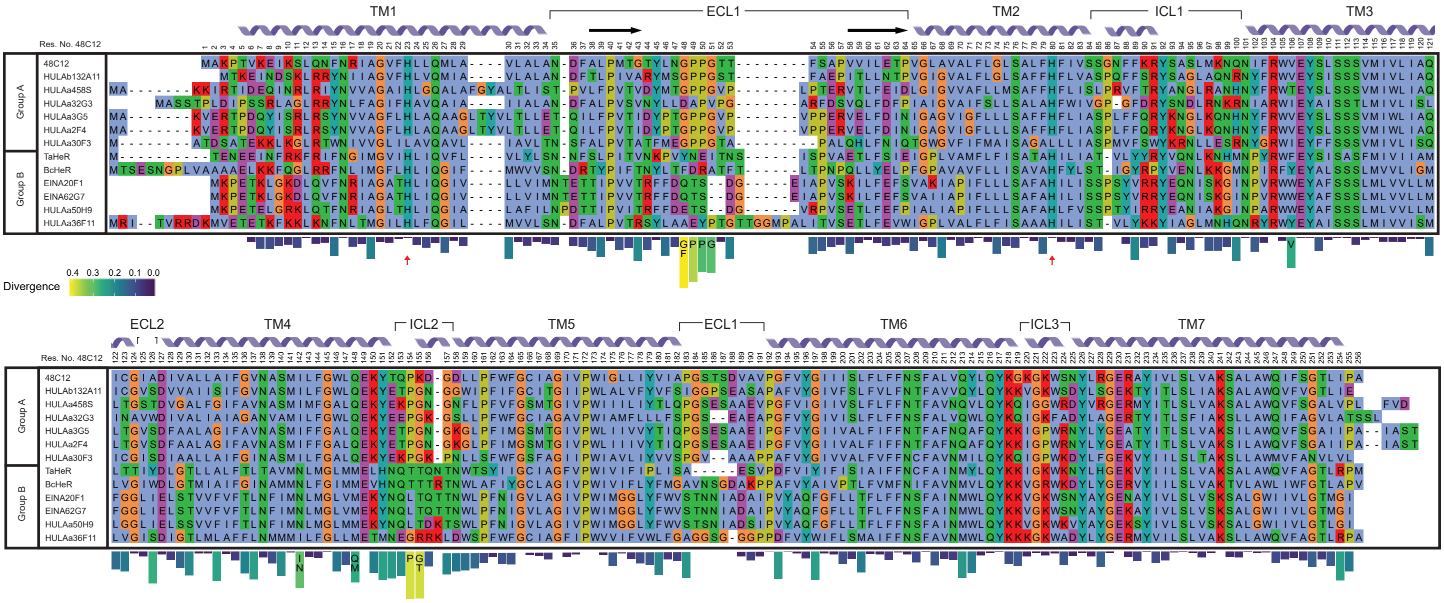


**Figure S5**. **Sequence alignment of HeRs from the environmental clones and previously characterized prokaryotic HeRs.** Part of a bigger alignment of prokaryotic HeRs from two phylogenetic groups (see Figure 1B) representing HeRs from the environmental clones from Ha'Hula and Ein Afek and three HeRs characterized previously. Structural annotation above the annotation corresponds to the structure of HeR-48C12 (PDB: 6SU3). Below the alignment a differential weblogo is provided showing Jensen-Shannon divergence per position between phylogenetic groups A and B, based on the alignment of 60%-identity clusters, with dashed positions representing alignment gaps. For positions with a divergence above 0.25, >50%-consensus residues are shown. Red arrows indicate the two highly conserved His residues that are mutated in HeR HULAa30F3.

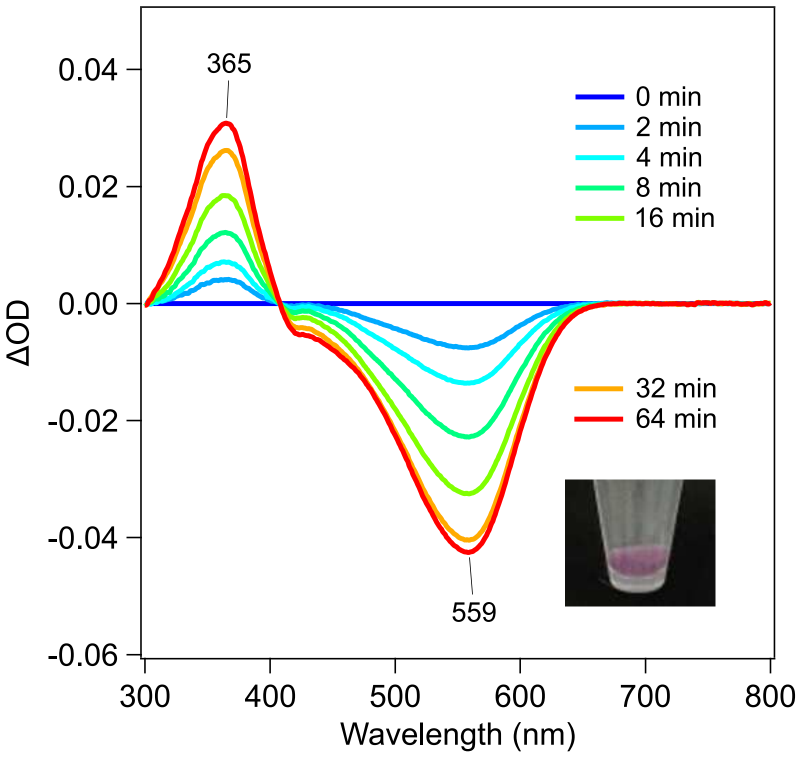


**Figure S6. Biophysical characterization of HeR HULAa30F3.** The difference in UV-vis absorption spectra of HeR HULAa30F3 between the spectra measured after and before bleaching of protein with hydroxylamine. The inset shows *E. coli* cell pellet expressing HeR HULAa30F3 in the presence of all-*trans* retinal.

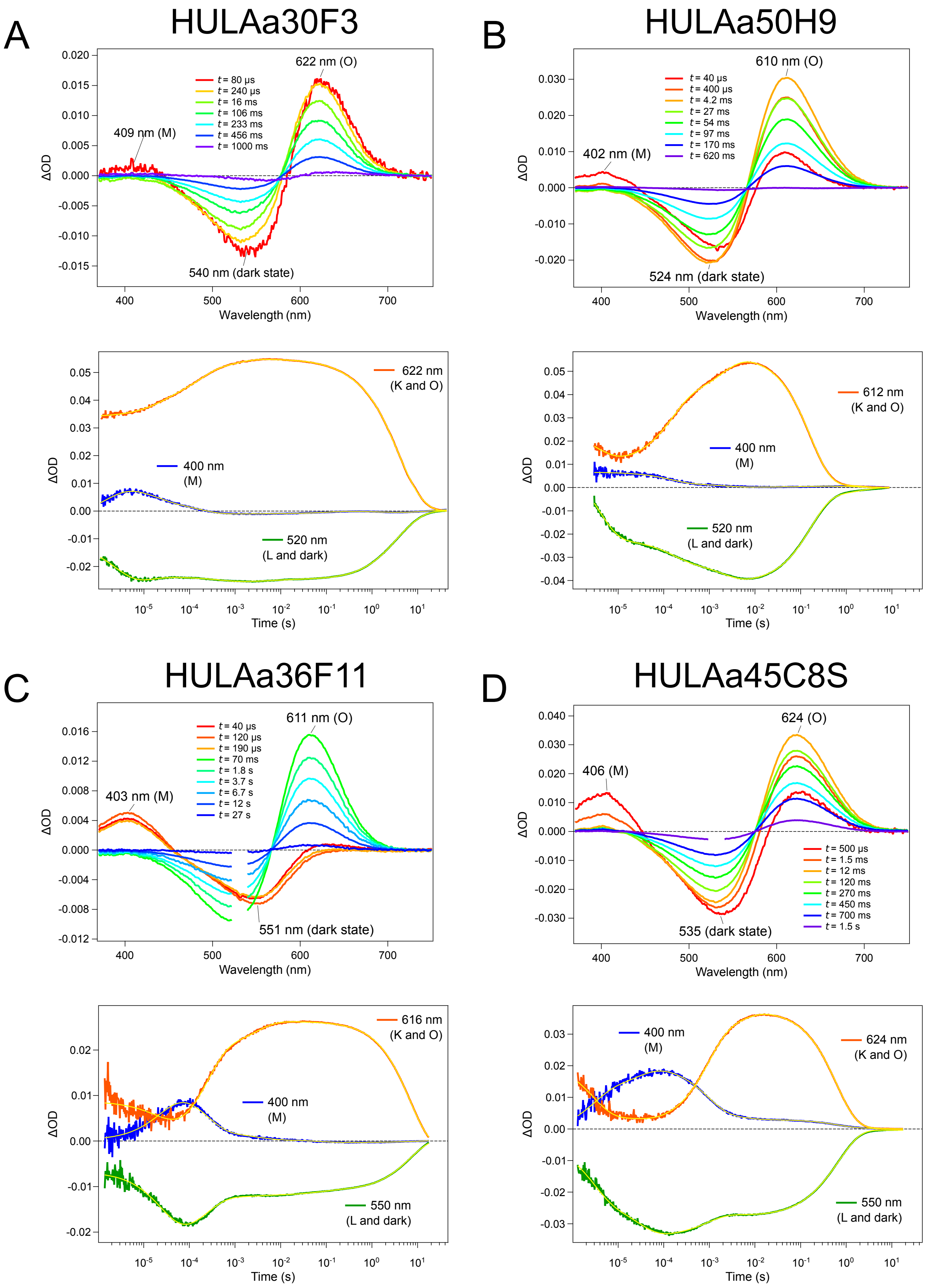
A
HULAa30F3
622 nm (O)
409 nm (M)
540 nm (dark state)
t = 80 µs
t = 240 µs
t = 16 ms
t = 106 ms
t = 233 ms
t = 456 ms
t = 1000 ms
ΔOD
Wavelength (nm)
622 nm
(K and O)
400 nm
(M)
520 nm
(L and dark)
Time (s)
B
HULAa50H9
610 nm (O)
402 nm (M)
524 nm (dark state)
t = 40 µs
t = 400 µs
t = 4.2 ms
t = 27 ms
t = 54 ms
t = 97 ms
t = 170 ms
t = 620 ms
612 nm
(K and O)
400 nm
(M)
520 nm
(L and dark)
C
HULAa36F11
611 nm (O)
403 nm (M)
551 nm (dark state)
t = 40 µs
t = 120 µs
t = 190 µs
t = 70 ms
t = 1.8 s
t = 3.7 s
t = 6.7 s
t = 12 s
t = 27 s
616 nm
(K and O)
400 nm
(M)
550 nm
(L and dark)
D
HULAa45C8S
624 (O)
406 (M)
535 (dark state)
t = 500 µs
t = 1.5 ms
t = 12 ms
t = 120 ms
t = 270 ms
t = 450 ms
t = 700 ms
t = 1.5 s
400 nm
(M)
624 nm
(K and O)
550 nm
(L and dark)

**Figure S7. Transient absorption changes of HeRs.** Transient absorption spectra (upper) and time evolutions of transient absorption change at specific wavelengths (lower), of HeR HULAa30F3 (**A**), HULAa50H9 (**B**), HULAa36F11 (**C**), and HULAa45C8S (**D**). The time evolutions of the accumulation of the K/O and the M, and the bleaching of the initial state were monitored at the wavelength indicated with orange (λ = 611-624 nm), blue (λ = 400-406 nm) and green (λ = 520-550 nm) solid lines. Yellow lines indicate the best-fit curves of multi-exponential function. For the measurement of transient spectra at $t$ > 70 ms and $t$ > 1 s for HULAa36F11 and HULAa45C8S, respectively, the jitter between laser pulse illumination and the exposure of detector could not be so precisely controlled due to instrumental limitation (see Materials and Methods), so that we needed to place a notch filter in front of a detector to avoid the saturation by scattered laser pulse. Hence, the absorption change could not be measured at $\lambda$ = 523-540 nm.

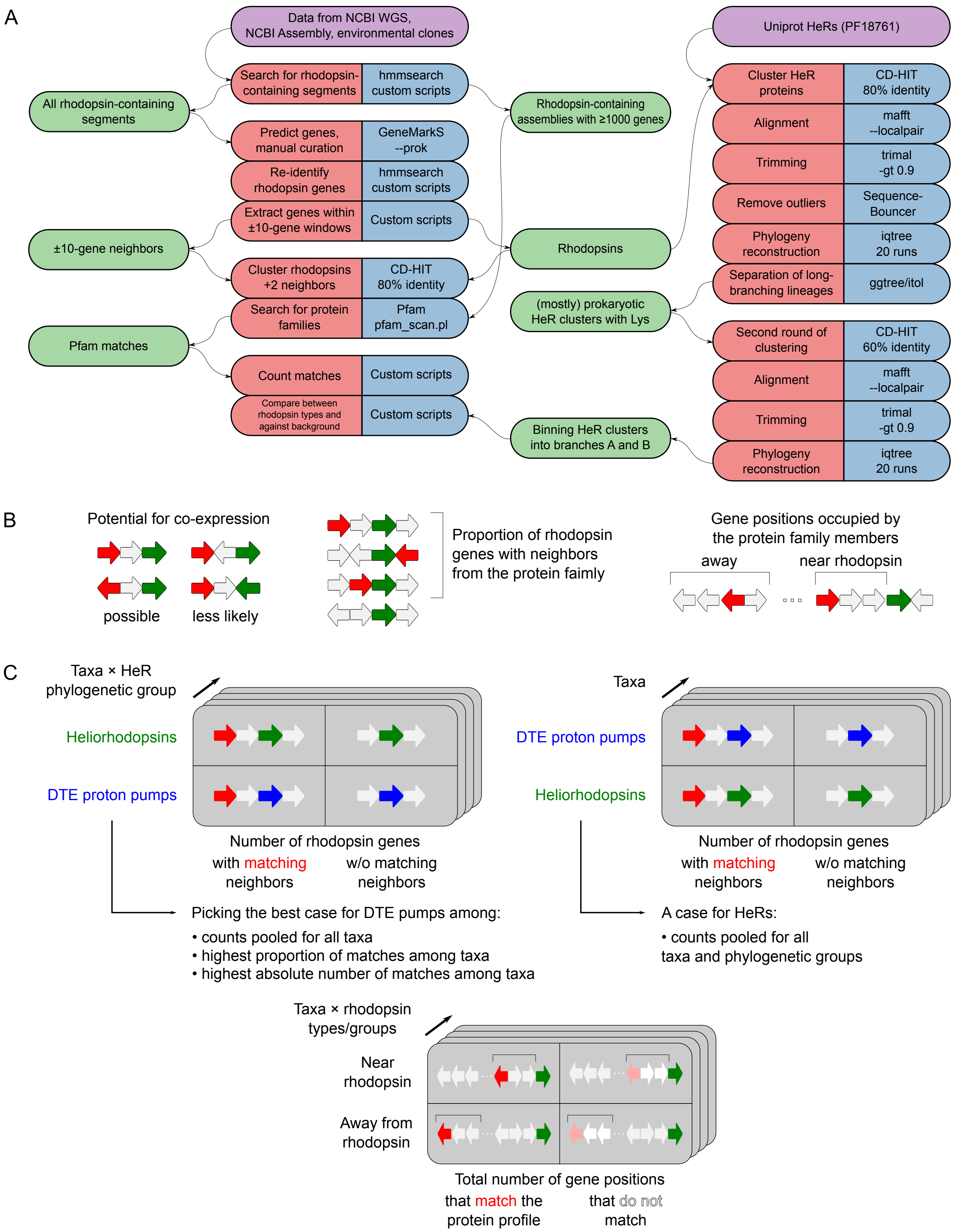
A
Data from NCBI WGS, NCBI Assembly, environmental clones
Search for rhodopsin-containing segments
hmmsearch custom scripts
All rhodopsin-containing segments
Rhodopsin-containing assemblies with ≥1000 genes
Predict genes, manual curation
GeneMarkS --prok
Re-identify rhodopsin genes
hmmsearch custom scripts
Extract genes within ±10-gene windows
Custom scripts
±10-gene neighbors
Rhodopsins
Cluster rhodopsins +2 neighbors
CD-HIT 80% identity
Search for protein families
Pfam pfam_scan.pl
Pfam matches
Count matches
Custom scripts
Compare between rhodopsin types and against background
Custom scripts
Binning HeR clusters into branches A and B
(mostly) prokaryotic HeR clusters with Lys
Uniprot HeRs (PF18761)
Cluster HeR proteins
CD-HIT 80% identity
Alignment
mafft --localpair
Trimming
trimal -gt 0.9
Remove outliers
Sequence-Bouncer
Phylogeny reconstruction
iqtree 20 runs
Separation of long-branching lineages
ggtree/itol
Second round of clustering
CD-HIT 60% identity
Alignment
mafft --localpair
Trimming
trimal -gt 0.9
Phylogeny reconstruction
iqtree 20 runs
B
Potential for co-expression
possible
less likely
Proportion of rhodopsin genes with neighbors from the protein faimly
Gene positions occupied by the protein family members
away
near rhodopsin
C
Taxa × HeR phylogenetic group
Heliorhodopsins
DTE proton pumps
Number of rhodopsin genes
with matching neighbors
w/o matching neighbors
Picking the best case for DTE pumps among:
• counts pooled for all taxa
• highest proportion of matches among taxa
• highest absolute number of matches among taxa
Taxa
DTE proton pumps
Heliorhodopsins
Number of rhodopsin genes
with matching neighbors
w/o matching neighbors
A case for HeRs:
• counts pooled for all taxa and phylogenetic groups
Taxa × rhodopsin types/groups
Near rhodopsin
Away from rhodopsin
Total number of gene positions
that match the protein profile
that do not match

**Figure S8. Pipeline used for phylogenetic classification of HeRs and for the analysis of neighbors. A**. Schematics of the different steps in the pipeline with the color code indicating: input data (lilac), computational step (red) with the corresponding software (blue) and derived data (green). **B**. Parameters estimated for each of the protein families found in the vicinity of rhodopsin genes: incidence of the gene orientations, proportion of HeR genes with the corresponding neighbors, proportion of gene positions occupied by the matching genes among rhodopsin neighbors and away from them (for long assemblies). **C**. Contingency tables created for the formal statistical tests for three types of comparisons: HeR for each grouping compared to the best case built for the DTE proton pumps; DTE proton pumps for each of the taxa against pooled counts for HeR neighbors; comparison of gene positions occupied by the corresponding neighbors against the background (gene positions away from the rhodopsin genes).

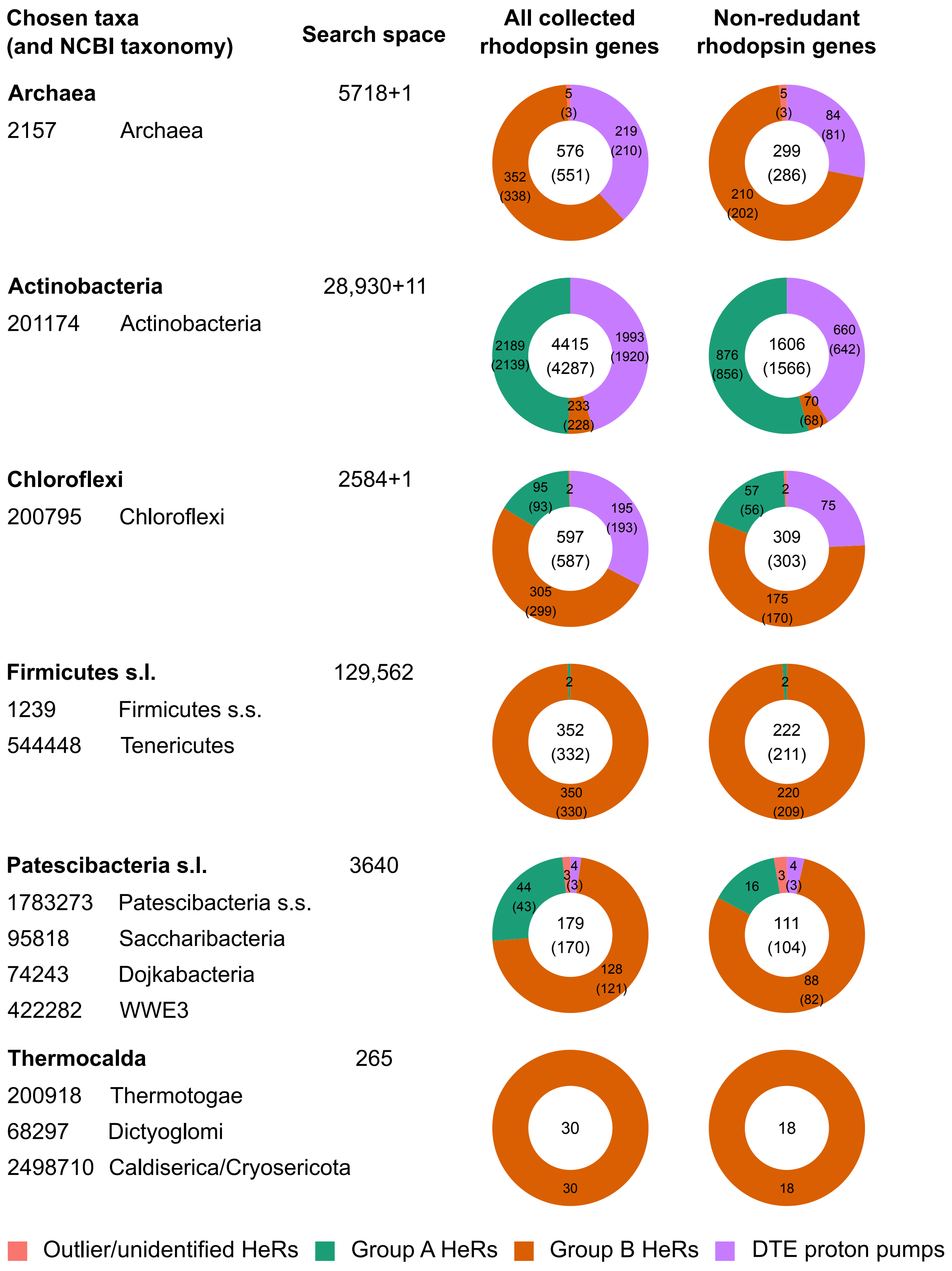
Chosen taxa
(and NCBI taxonomy)
Search space
All collected
rhodopsin genes
Non-redudant
rhodopsin genes
Archaea
5718+1
2157 Archaea
5
(3)
219
(210)
576
(551)
352
(338)
5
(3)
84
(81)
299
(286)
210
(202)
Actinobacteria
28,930+11
201174 Actinobacteria
2189
(2139)
4415
(4287)
1993
(1920)
233
(228)
876
(856)
1606
(1566)
660
(642)
70
(68)
Chloroflexi
2584+1
200795 Chloroflexi
95
(93)
2
195
(193)
597
(587)
305
(299)
57
(56)
2
75
309
(303)
175
(170)
Firmicutes s.l.
129,562
1239 Firmicutes s.s.
544448 Tenericutes
2
352
(332)
350
(330)
2
222
(211)
220
(209)
Patescibacteria s.l.
3640
1783273 Patescibacteria s.s.
95818 Saccharibacteria
74243 Dojkabacteria
422282 WWE3
44
(43)
3
4
(3)
179
(170)
128
(121)
16
3
4
(3)
111
(104)
88
(82)
Thermocalda
265
200918 Thermotogae
68297 Dictyoglomi
2498710 Caldiserica/Cryosericota
30
30
18
18
Outlier/unidentified HeRs
Group A HeRs
Group B HeRs
DTE proton pumps

**Figure S9. Taxa used for the analysis of rhodopsin neighbors.** Summary statistics for the six taxa chosen for the analysis of HeR neighbors. The taxa were chosen to represent a sufficient number of phylogenetically close and morphologically similar prokaryotes and represent the vast majority of HeR-harboring prokaryotes based on the selection of HeR proteins in Uniprot. The “search space” indicates the number of the analyzed NCBI WGS and Assembly records assigned to the corresponding NCBI taxonomy accessions (the number after “+” indicates the number of HeR-coding environmental clones). The name `Patescibacteria` is used here in the broader sense to include all “candidate phyla” of which the four biggest groups of HeR-harboring assemblies were analyzed. Thermocalda unites three phyla or classes of diderms lacking lipopolysaccharides in the outer membranes following. The total number of the analyzed assemblies, raw number of the rhodopsin genes with neighbors identified (according to the filtering criteria – see Materials & Methods) and the numbers of non-redundant rhodopsin genes (i.e. having distinct triads: rhodopsin and two of its immediate neighbors, see Supplementary Figure S8) are provided. In parenthesis are the numbers of assemblies for each category (if different from the number of the genes, the discrepancy arises from assemblies containing multiple HeR genes). HeR are subdivided into the phylogenetic groups A and B (see Figures 1 and Supplementary Figure S5).

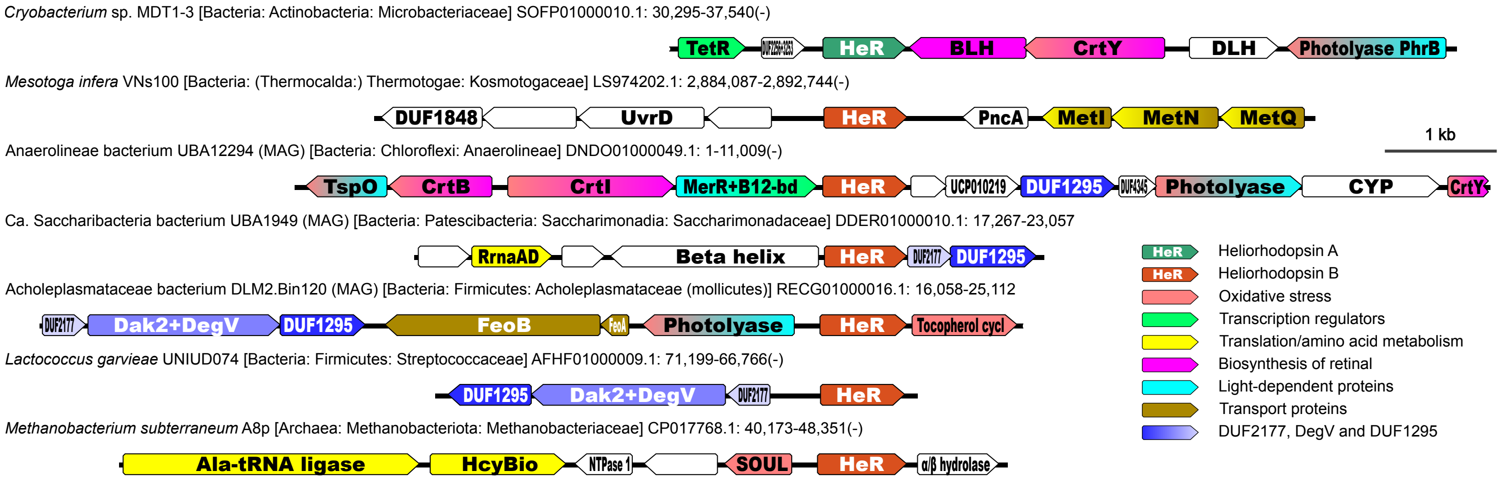


**Figure S10. Examples of HeR gene neighbors in various prokaryotes.** Representative DNA fragments containing HeR genes from different prokaryotic groups with the neighboring genes classified into functional categories. Note that some gene products are classified in two categories and receive gradient color. The examples include genome and metagenome assemblies of: *Cryobacterium* sp. MDT1-3 with a rare case of a HeR gene next to *blh*; *Mesotoga infera* as a representative of unusual diderms; Anaerolineae bacterium UBA12294, a metagenomic assembly in which the HeR gene is surrounded by several genes of the beta-carotene biosynthetic pathway but not *blh*, as well as a putative cobalamine-binding (“B12-bd”) transcription regulator; Acholeplasmataceae bacterium DLM2.Bin120, a mollicute with a tocopherol cyclase gene typical for this group, as well as the triad DUF2177, DegV and DUF1295; *Lactococcus garvieae*, a lactic bacterium with the triad DUF2177, DegV and DUF1295; *Methanobacterium subterraneum* A8p, a methanogenic archaeon with a gene coding for a SOUL-domain-containing heme-binding protein and multiple genes involved in translation/amino acid metabolism.

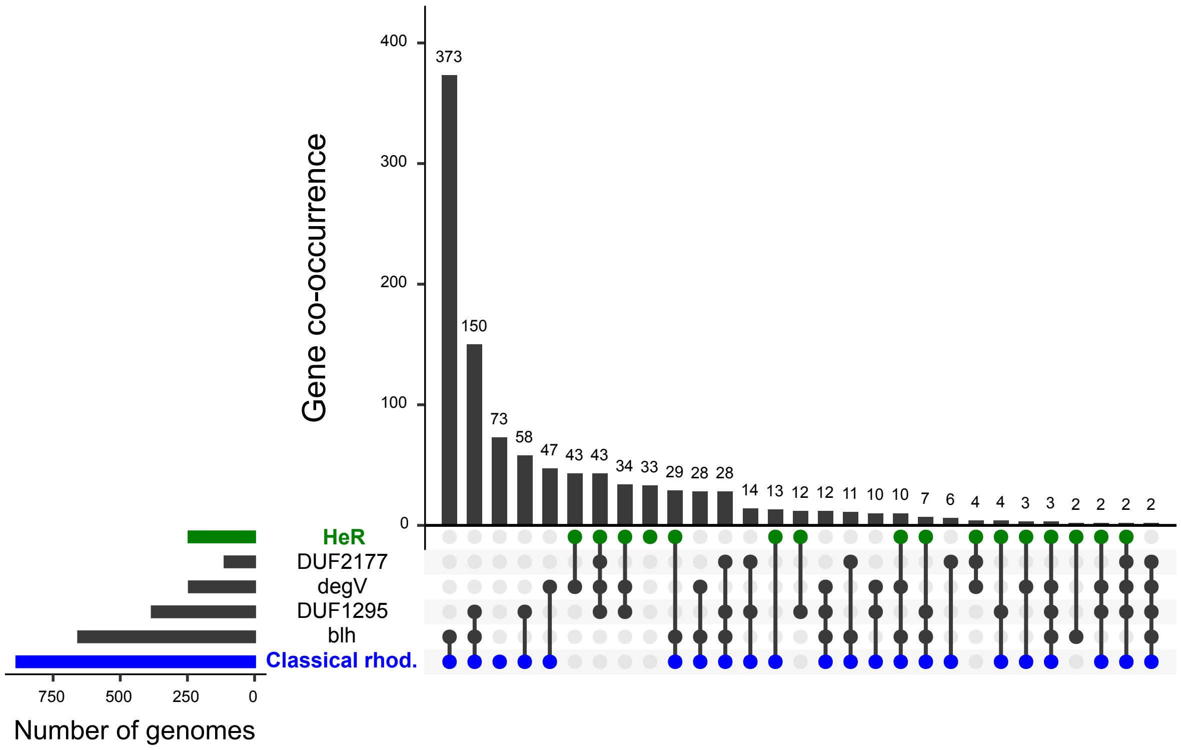


**Figure S11. Genome-wide co-occurrences of protein families characteristic to rhodopsin neighbors.** Co-occurrence of HeRs (PF18761) and other rhodopsin types (PF01036) with four chosen protein families frequently co-locating with them. The search covered 18,708 of high-quality non-redundant prokaryotic assemblies among GTDB representative genomes. Note that the protein families DUF2177, DegV and DUF1295 alone have a much wider distribution than the rhodopsin genes, therefore only gene combinations involving rhodopsins are shown.
